# A human epithelial co-culture system reveals distinct host cell interaction behaviours for *Treponema pallidum*

**DOI:** 10.64898/2026.02.04.703719

**Authors:** Katia Capuccini, Divenita Govender, David Goulding, Cecilia Kyanya, Shivani Pasricha, Lorenzo Giacani, Nicholas R Thomson, Linda Grillová

## Abstract

*Treponema pallidum* subsp. *pallidum*, the causative agent of syphilis, remains difficult to study owing to long-standing limitations in *in vitro* cultivation. Although a rabbit epithelial cell co-culture system (Sf1Ep) enabled major advances, the absence of human cell-based models limits clinical relevance and mechanistic understanding of host–pathogen interactions. Here, we established a co-culture system using human epithelial cell lines capable of supporting *T. pallidum* growth *in vitro*. Six epithelial or epithelial-like cell lines from diverse tissues were evaluated under microaerophilic conditions using standard cultivation medium. CAL-39 (vulva) and HepG2 (liver) supported *T. pallidum* survival, replication, characteristic growth behaviors, and long-term passage at levels comparable to Sf1Ep cells. Growth kinetics, attachment dynamics, and motility were quantified over extended culture. Using live-cell imaging, we defined two distinct host-cell interaction behaviors: surface-associated crawling and stable single-polar attachment. Both behaviors were observed across all tested cell lines but persisted only in those permissive for sustained growth. Together, our findings establish clinically relevant human epithelial co-culture models for *T. pallidum*, provide new insights into host-cell-dependent growth and motility, and create a platform for mechanistic studies of syphilis pathogenesis and vaccine target discovery.

## Introduction

Syphilis, caused by *Treponema pallidum* subsp. *pallidum* (*T. pallidum*), typically begins with a localized chancre at the site of infection, and rapidly disseminates in multiple tissues and organs, leading to a wide spectrum of clinical manifestations across the primary, secondary, and tertiary stages. If left untreated, syphilis can result in severe and irreversible neurological and cardiovascular complications and remains a leading cause of adverse pregnancy outcomes worldwide, including stillbirths and neonatal deaths when congenital infection occurs (1). Although syphilis is readily curable with appropriate antibiotic therapy, its incidence has increased substantially over the past two decades, particularly among specific communities, including men who have sex with men. The WHO estimates approximately 8 million new syphilis infections worldwide in 2022, compared to 7.1 million cases in 2020 (2). Together, the rising incidence and burden of preventable morbidity and mortality highlight the urgent need to strengthen syphilis prevention, screening, and control strategies through a deeper understanding of basic biology as well as testing.

While the development of an effective vaccine would significantly help to control the spread of syphilis, there is currently little consensus on target selection. Importantly, while there is a strong focus on syphilis vaccine development, progress is hindered by a lack of understanding of the pathogenesis and basic biology of *T. pallidum*. For decades investigators have been unable to steadily propagate this bacterium *in vitro*, needing to rely on *in vivo* propagation in the rabbit infection model.

However, this changed in 2018, when a system for successful long-term *in vitro* co-cultivation of *T. pallidum* with rabbit skin epithelial cells (Sf1Ep) was established (3). This breakthrough accelerated progress in the field, with significant advancements including the development of methods for genetic modification of *T. pallidum* (4), high-throughput antibiotic sensitivity/resistance testing (5), and proteomics and transcriptomic studies (6, 7), among other tools and procedures that are standard in traditional microbiology for different bacteria.

The Sf1Ep co-culture system is valuable for propagating *T. pallidum in vitro*. However, to establish accessible and clinically relevant models of early host–pathogen interactions at epithelial surfaces, where the bacterium first encounters the human host, there is a need for infection systems based on human rather than rabbit cells. Because interactions with the pathogen are likely to vary between cell types and species, we sought to establish more clinically relevant *in vitro* infection models by identifying suitable human cell lines that could replace rabbit epithelial Sf1Ep cells as a foundation for future mechanistic studies.

We identified two human epithelial cell lines from distinct body sites – vulva and liver – that support sustained growth and the complete growth cycle of *T. pallidum* in a manner comparable to Sf1Ep cells. For the first time, by combining these novel, clinically relevant co-culture models with advanced microscopy platforms, we describe two distinct modes of behaviour during *T. pallidum* interaction with human and rabbit epithelia. These consist of both surface-associated crawling and stable, single-polar attachment, observed using live imaging of replicating cultures, as well as in fixed samples. We anticipate that human cell models that support *T. pallidum* growth will enable a new generation of clinically-grounded mechanistic discoveries, accelerating vaccine target discovery and closing critical knowledge gaps in *T. pallidum*-host interactions. These insights will ultimately transform control of syphilis, a disease that remains poorly understood despite its long history.

## Results

### Selection of human cell lines

Currently, *T. pallidum* is cultured using Sf1Ep rabbit epithelial cells under low-oxygen conditions (1.5% O2) in *T. pallidum* cultivation medium (TPCM-2) for 7 days, reflecting its long generation time (>35 h) (3). To identify human alternatives to rabbit cells, we selected six well-characterized human epithelial or epithelial-like cell lines prioritising hypoxia-tolerant cell lines with longer doubling times (>30 h), replicating key characteristics of Sf1Ep cells, while including a faster-growing cell line (HEK-293, <20 h) as a comparator (Table 1). The selected cell lines span diverse tissue origins: vulva (CAL-39, SW954), placenta (JAR, JEG-3), liver (HepG2), and kidney (HEK-293). All cell lines were first assessed for growth in 1.5% O2 and in TPCM-2 over a 7-day period, replicating the established Sf1Ep co-culture workflow (3). While some cell lines exhibited longer doubling times under these conditions compared to optimal normoxic conditions typical of human cell culture (Table 1), none showed morphological abnormalities or signs of significant cell damage when cultured under low oxygen (see Method section for details).

**Table 1.**
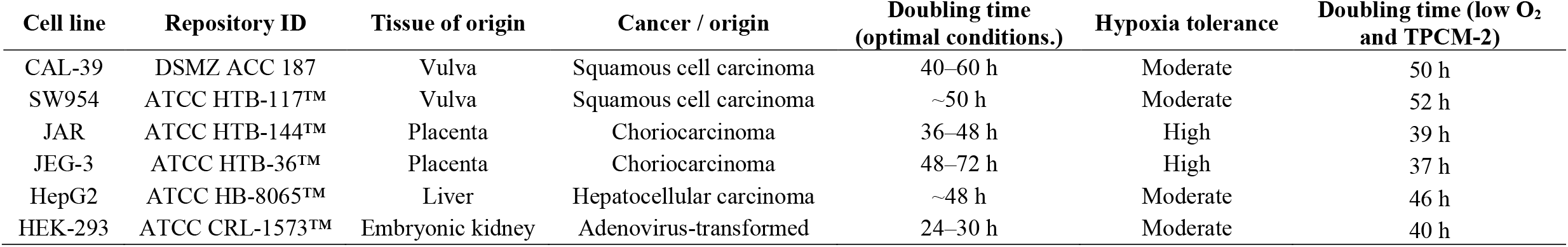
Human cell lines tested for *T. pallidum* growth support.

### Testing human cell lines for T. pallidum in vitro growth support

We first assessed *T. pallidum* growth using the SS14 strain across the six selected human cell lines and compared these to *T. pallidum* cultured without mammalian cells (negative control, NC) or cocultured with the reference supporting cell line Sf1Ep (positive control, PC), over a 4-week period (Fig. 1A). At each weekly passage, cultures were reseeded with a fixed inoculum of 2.5 × 10^6^ *T. pallidum* cells that attached to the host cell monolayer. We observed that both subpopulations of *T. pallidum* cells - those attached to the mammalian cell surface and the non-attached cells in a planktonic state in the medium - remain viable under standard culture conditions. This was shown by re-seeding experiments in which *T. pallidum* cells collected exclusively from the culture medium reattached to fresh monolayers and completed a full growth cycle (Supplementary Fig. 1). Therefore, both attached and non-attached *T. pallidum* populations were assessed during subsequent characterisation of growth dynamics in this study. The only exception was this specific experiment, in which, apart from a negative control, only monolayer-attached *T. pallidum* cells were used to estimate growth kinetics, in accordance with standard *T. pallidum* passage recommendations (8). For the HEK-293 and JAR cell lines, *T. pallidum* cell numbers fell below this threshold during the first and third passages, respectively, and the cultures were terminated after weeks 1 and 3. As expected, *T. pallidum* negative control cultures showed markedly reduced total *T. pallidum* yields over time. Among the six candidate cell lines tested, CAL-39 and HepG2 displayed *T. pallidum* growth profiles that were comparable to the Sf1Ep standard (Fig.1A). Comparison of total *T. pallidum* counts using Tukey’s HSD test showed no significant difference in total *T. pallidum* counts between *T. pallidum* co-cultured with CAL-39 or HepG2 and *T. pallidum* co-cultured with Sf1Ep (p > 0.05). By contrast, *T. pallidum* co-cultured with either CAL-39 or HepG2 cells showed significantly higher total *T. pallidum* counts compared to *T. pallidum* NC (p < 0.05; see Supplementary Fig. 2 for p-values for all pairwise comparisons across cell lines).

**Figure 1.**
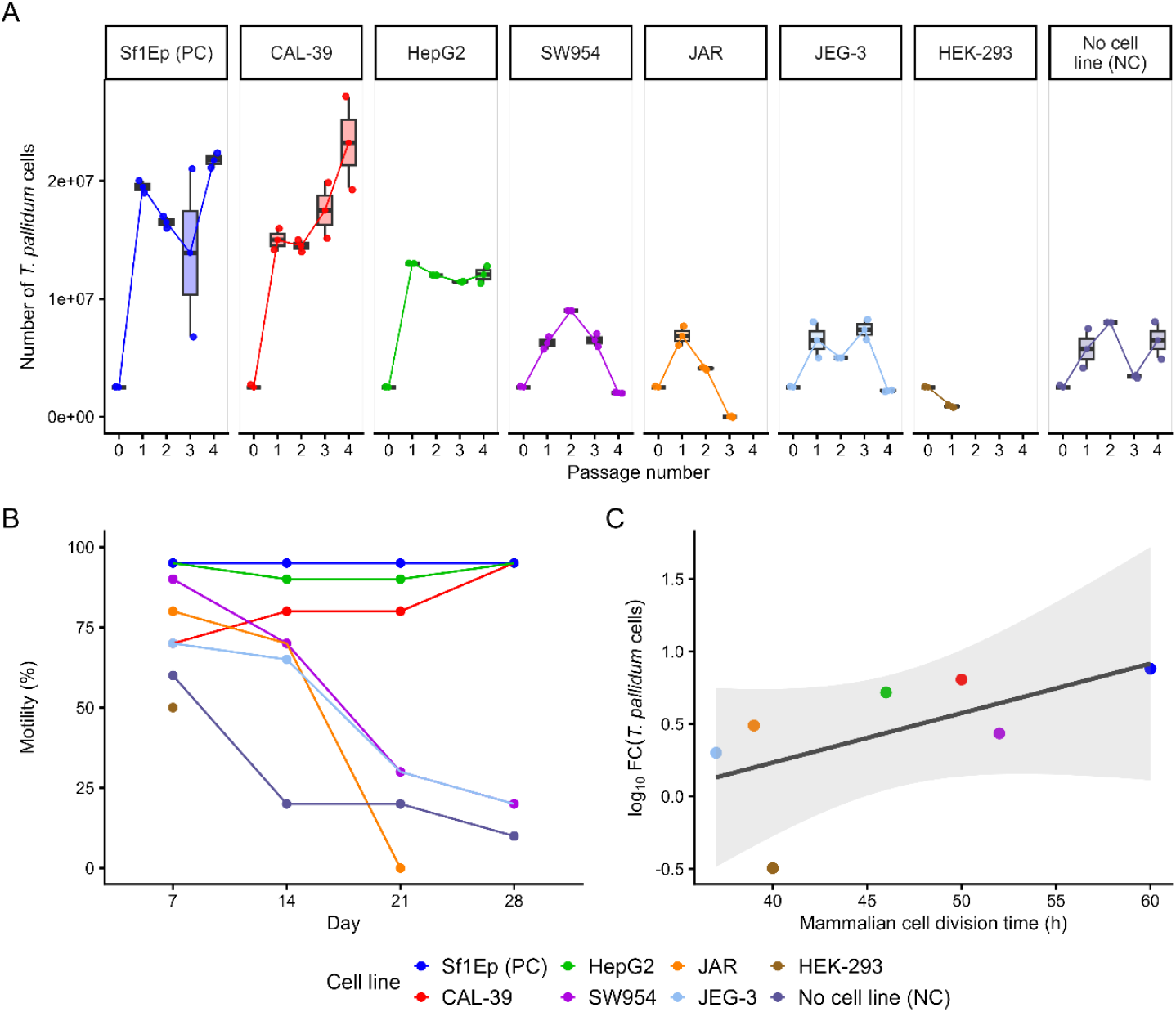
Human epithelial cell line–dependent growth, motility, and yield of *T. pallidum* in long-term co-culture. (A) *T. pallidum* growth in the presence of different human cell lines over 28 days. Connected boxplots depict *T. pallidum* cell counts from three biological replicates, measured at weekly harvests, with lines connecting mean yields across passages. For each weekly passage, a standardized treponemal inoculum (2.5 × 10^6^ cells) was used to subculture into wells freshly seeded with human cells. For HEK-293 (brown) and JAR (orange) cell lines, cultures were terminated early due to insufficient *T. pallidum* inoculum (<2.5 × 10^6^ cells) at passage 1 and passage 3, respectively. NC, negative control (*T. pallidum* cultured without host cells); PC, positive control (*T. pallidum* cultured with rabbit epithelial Sf1Ep cells). (B) Percentage of motile *T. pallidum* cells across co-cultures over 4 weeks. Points represent mean values from biological replicates. (C) Relationship between mammalian cell division time and *T. pallidum* yield. Scatterplot shows log_10_ *T. pallidum* cell number fold change versus mean mammalian cell division time (hours), with the solid line representing linear regression and shaded 95% confidence interval. Regression analysis indicated a positive but non-significant association (slope = 0.034; R^2^ = 0.37; p = 0.147).

Across all time points, *T. pallidum* motility remained high (∼90%) in CAL-39, HEPG2, and Sf1Ep, but was significantly reduced in the other tested cell lines (Fig. 1B; p = 0.0055). Motility did not differ significantly among CAL-39, HEPG2, and Sf1Ep. All cell lines increased motility relative to the no-cell control, reaching statistical significance for HEPG2 and Sf1Ep, while CAL-39 showed a consistent but non-significant increase, supporting its suitability as a co-culture model.

### Relationship between mammalian cells division time and T. pallidum yield

To explore the relationship between human cell division time and *T. pallidum* yield, *T. pallidum* fold change was plotted against the division time of each human cell line, with Sf1Ep cells included as a reference (Fig. 1C). *T. pallidum* yield, expressed as log_10_ fold change, increased with longer host cell division times in both human and rabbit cells (slope = 0.034 log_10_ units per hour). This relationship explained 37% of the variance but did not reach statistical significance (R^2^ = 0.37, p = 0.147).

### *Longitudinal T. pallidum growth dynamics* in *in vitro* co-culture

To characterize *T. pallidum* growth across the different host cell lines and to determine the timing of exponential growth of *T. pallidum*, cultures were monitored daily for 10-days. Each 24-hour time point was analysed using a dedicated culture plate, with all plates inoculated simultaneously on day 0 with the same starting inoculum; at each time point a single plate was terminated and processed for *T. pallidum* quantification. In this experiment, growth was estimated from the host cell attached as well as the non-attached *T. pallidum* cells floating in the culture medium to provide a more detailed view of *T. pallidum* growth dynamics.

Cell lines Sf1Ep, CAL-39, and HepG2 all supported similar *T. pallidum* growth trajectories, with comparable increases in total *T. pallidum* cell count over the 10-day period (Fig. 2A). The relative proportions of attached and planktonic *T. pallidum* populations in these cultures are shown in Fig. 2C. After day 9 of culture, the medium in CAL-39 and HepG2 co-cultures became acidified, as indicated by a colour change to yellow, and *T. pallidum* cell numbers in both attached and planktonic fractions began to decline. Intact and motile treponemes were still detectable by microscopy, but at lower abundance, suggesting that cell lysis may contribute to the reduced counts under these conditions. By contrast, no comparable decline was observed in Sf1Ep co-cultures. Therefore, currently culturing *T. pallidum* with these human cell lines would not be recommended beyond 9 days, while the standard 7-day culture period is recommended for consistency and convenience.

**Figure 2.**
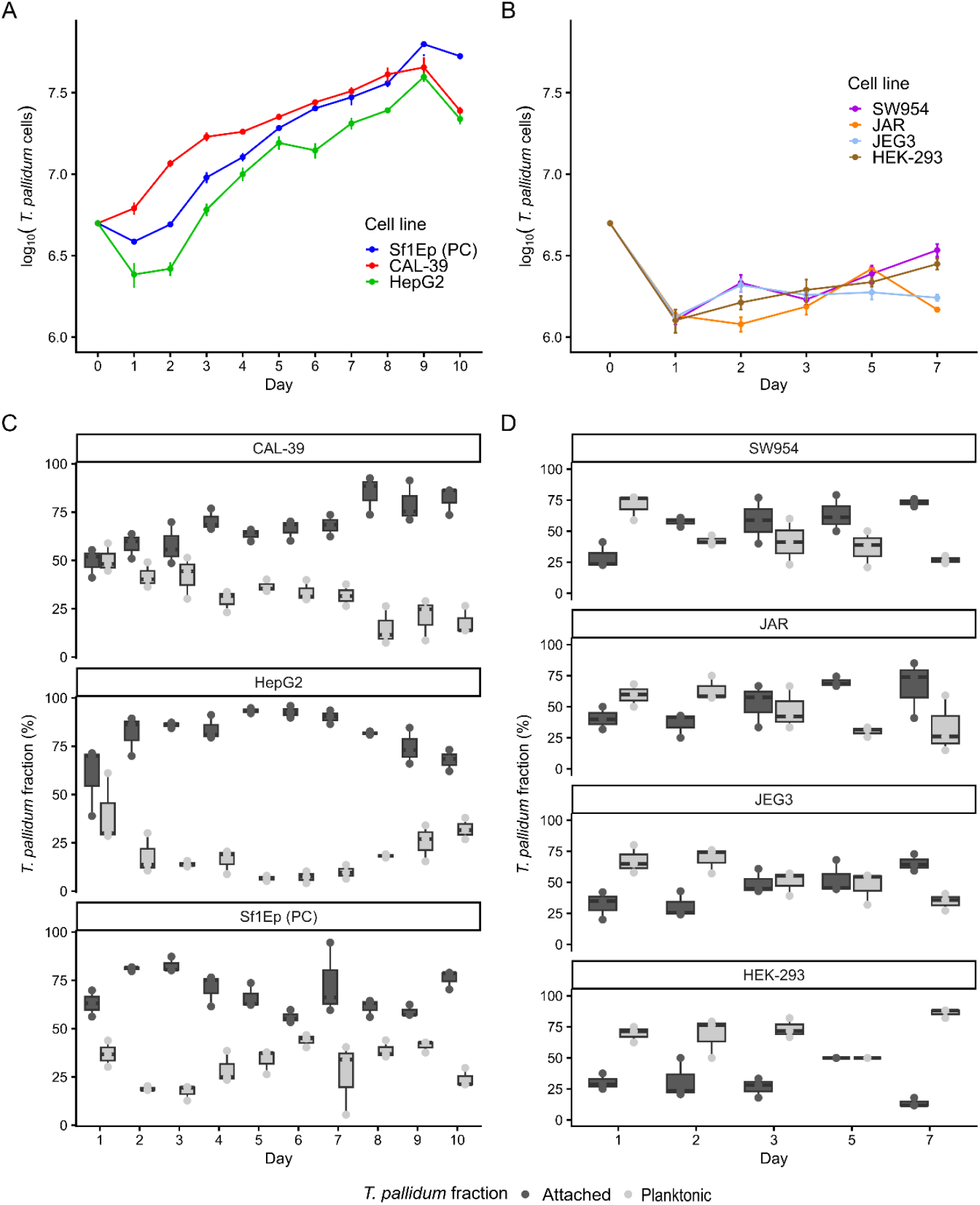
Growth of *T. pallidum* in supporting and non-supporting mammalian cell lines. (A) Log_10_ *T. pallidum* counts in supporting cell lines (Sf1Ep, CAL-39, and HEPG2). (B) Log_10_*T. pallidum* counts in non-supporting cell lines (HEK-293, SW954, JAR, JEG-3). (C) Boxplots showing percentages of attached (dark grey) and planktonic (light grey) fractions in supporting cell lines. (D) Boxplot showing percentages of attached (dark grey) and planktonic (light grey) fractions in non-supporting cell lines. Data points represent mean values across biological replicates. For supporting cell lines, there is no significant difference between conditions between day 1 and 7 (p-values for all comparisons for each day reported in Supplementary Table 1). For non-supporting cell lines, differences between cell lines were also not significant across the whole experiment (ANOVA p = 0.142).

In parallel, *T. pallidum* growth was monitored in the non-supporting cell lines HEK-293, SW954, JAR, and JEG-3 at days 1, 3, 5, and 7 (Fig. 2B). Total *T. pallidum* counts in these non-supporting cell lines were substantially lower than those observed in the supporting cell lines, with a maximum of 3.5 × 10^6^ cells. Moreover, a higher proportion of *T. pallidum* remained in the planktonic fraction when co-culturing with non-supporting cell lines, consistent with reduced attachment to the host-cell monolayer (Fig. 2D). Wilcoxon rank-sum tests conducted independently for each day yielded p-values of 0.057 (day 1), 0.057 (day 2), 0.114 (day 3), 0.229 (day 5), and 0.229 (day 7). Corresponding boxplots and statistical comparisons are presented in (Supplementary Fig. 3).

### Real time interaction of T. pallidum with host cell lines

To visualise *T. pallidum* interactions with host cells in real time, we established the first live-cell imaging platform for *T. pallidum*, enabling continuous monitoring of bacterial behaviour during co-culture. This approach was first used to characterise host–pathogen interactions in the standard Sf1Ep co-culture system. Two distinct populations of *T. pallidum* interacting differently with host cells were identified. First, *T. pallidum* exhibited highly dynamic, surface-associated motility characterized by close contact with host cell membranes and movement along the cellular surface (Video file, Fig. 3A). This behaviour resembles the “crawling” movement, or “translocation via an undulating waveform,” previously described for *T. pallidum* interactions with human platelets (9) and for other pathogenic spirochetes such as *Leptospira* and *Borrelia* (10, 11). *T. pallidum* cells displayed back-and-forth (reversal) movement using alternating poles and, in some cases, turned in the opposite direction using the same pole, likely due to transient self-entanglement (Video file, Fig. 3B). Second, a distinct population of *T. pallidum* cells exhibited stable single-polar attachment to the host cell surface, with the opposite pole remaining free (Video file, Fig. 3C).

**Figure 3.**
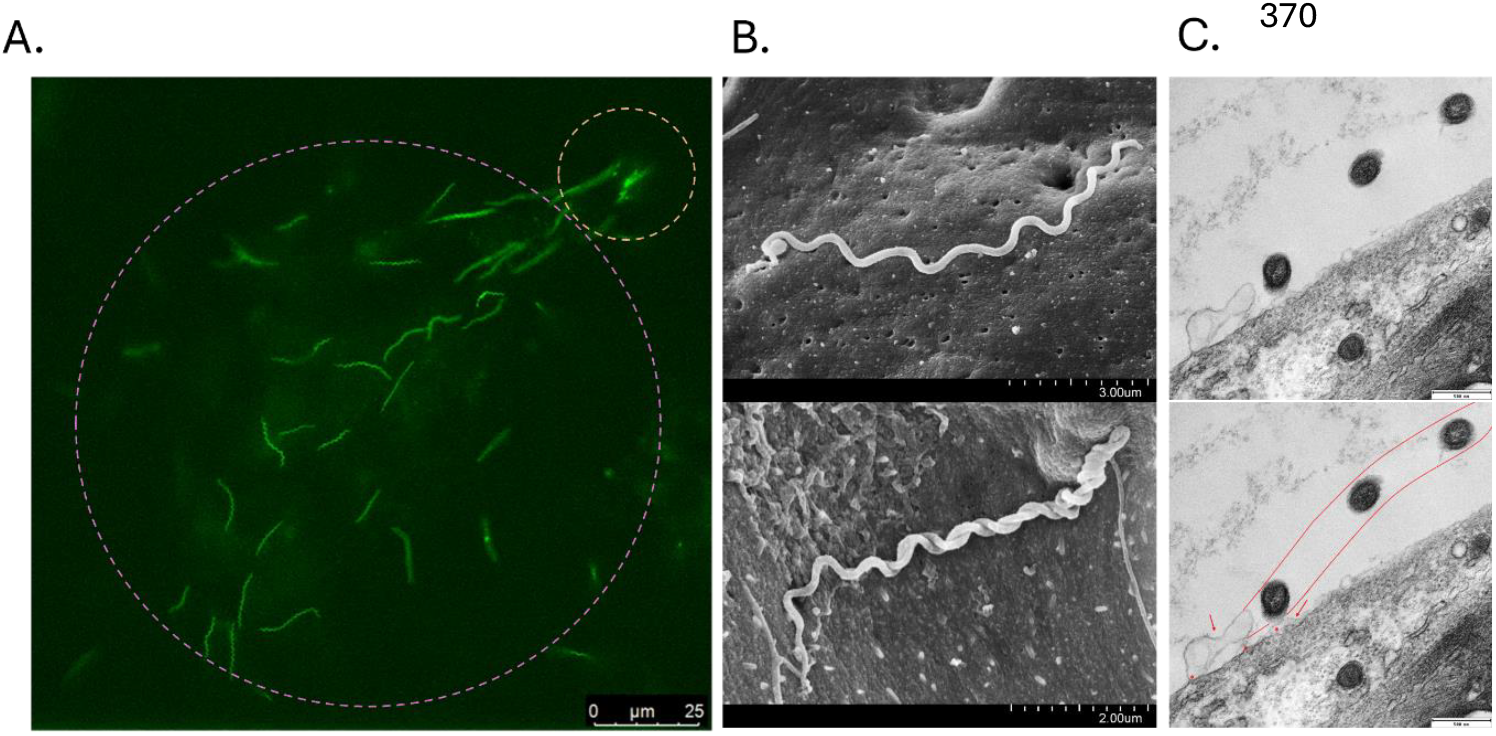
Interaction of *T. pallidum* with host cell lines. (A) Snapshot from live-cell imaging (Video file) showing two populations of *T. pallidum* interacting with the host cell surface. The population marked by the circle in the top right represents cells exhibiting single polar attachment, while the population marked by the circle in the bottom left represents *T. pallidum* cells displaying crawling-like motility. (B) SEM of crawling-like *T. pallidum* populations. The top panel shows a *T. pallidum* cell fully extended on the host cell surface, remaining attached along its length while moving across the cell. The bottom panel shows a *T. pallidum* cell captured at the moment of turning in the opposite direction, also observed in the Video file. (C) TEM of *T. pallidum* cells attached to the host cell surface by a single pole. The red line indicates the localization of *T. pallidum* cell, and the stars and arrows indicate possible attachment sites.

Next, real-time imaging of *T. pallidum* interacting with all tested host cell lines was performed, and these interactions were monitored over a 10-day period to complement the full picture of *T. pallidum* growth dynamics. Interestingly, the characteristic behaviours described in this study, represented by two motility types – crawling-like and single–pole–attached – were observed in all host cell lines tested during the first 3 days (including rabbit Sf1Ep cells and human Cal-39, HepG2, JEG-3, JAR, HEK-293, and SW954 cells). In Sf1Ep, Cal-39, and HepG2 cells, both motility types persisted throughout the full 10 days of observation. By contrast, in human cell lines described as non-supportive for *T. pallidum* growth (JEG-3, JAR, HEK-293, and SW954), *T. pallidum* cells exhibiting crawling-like behaviour gradually disappeared; overall, cells became increasingly non-motile, with the only remaining motile cells showing single-polar attachment.

## Discussion

Since 2018, *T. pallidum* is cultured *in vitro* using Sf1Ep rabbit epithelial cells in *T. pallidum* cultivation medium (TPCM-2; media (3)) and under low-oxygen conditions (1,5% O_2_) for 7 days, a duration required to allow *T. pallidum* multiplication due to its long generation time (more than 35 hours). Over the past several decades, numerous culture media formulations and multiple cell lines were tested, but only the combination of TPCM-2 and Sf1Ep cells proved effective (3). Historically tested human, rabbit, and rat cell types included those from testis, kidney, spleen, lung, epidermis, cervix, urethra, and nerve tissues. Although *T. pallidum* showed affinity for these cells, it did not survive long term, persisting for no more than a few days. (12). Importantly, the human cell lines tested were predominantly fast-growing (with doubling time less than 20 hours in optimal conditions), in contrast to the much slower-growing Sf1Ep cells (with doubling time more than 30 hours in optimal conditions). Moreover, using higher passage (>60) Sf1Ep cells in the co-culture system results in a decreased treponemal yield and motility (8). This is likely because high-passage Sf1Ep cells tend to divide more rapidly. Similarly, starting the culture with more than 10^5^ Sf1Ep cells per well of a 6-well plate have been found to result in lower culture yield and decreased treponemal motility (8). Therefore, a high density of Sf1Ep cells appears to negatively affect treponemal growth, likely due to competition between Sf1Ep and treponemes for shared nutrients, accumulation of secondary metabolites, and consequent acidification of the culture medium.

Given the above, we hypothesised that slow-growing human cell lines with longer generation times may have greater potential to support a slow-growing bacterium such as *T. pallidum*. Hence, our human cell line selection was guided by the premise that reduced host-cell proliferation may limit nutrient competition and media acidification, and that adaptation to low-oxygen environments may make certain human cell lines more permissive under the hypoxic conditions (1.5% O2) required for *T. pallidum* cultivation. Among the six host cell lines tested, CAL-39 and HepG2 supported *T. pallidum* growth at levels comparable to the reference Sf1Ep cell line. In contrast, but consistent with previous studies, *T. pallidum* cultured without any eukaryotic cell showed markedly reduced total cell counts over time, consistent with a stress condition leading to progressive cell loss (13). A similar outcome was observed with non-supporting cell lines, in which *T. pallidum* cell numbers frequently declined below the threshold required for continued passaging, indicating that a specific host–cell interaction is necessary to sustain *T. pallidum* growth *in vitro*. Importantly, although human cell division time may contribute to cell permissiveness, regression analysis in this study showed only a positive but non-significant association (Figure 1C). This suggests it is unlikely to be the primary determinant, and that other factors, such as media composition, surface receptor expression, metabolic profiles or stress-response pathways of human cells, may also play a role.

The establishment of a live-cell imaging platform enabled, for the first time, direct visualisation of *T. pallidum* interactions with host cells under physiologically relevant, microaerophilic conditions. It showed that *T. pallidum* can initially colonize all tested cell lines, regardless of their ability to support long-term growth, at least during the first 3 days of co-culture. This observation is consistent with extensive clinical and historical evidence demonstrating dissemination of *T. pallidum* to a wide range of human tissues in atypical and systemic syphilis, including bone (14), gastric mucosa (15), liver (16), extragenital derma (17), lungs (18), thyroid gland (19), eyes (20) and even multiple sites simultaneously (21). However, *in vitro, T. pallidum* is likely to engage in distinct molecular interactions with host cells that support sustained bacterial growth compared with cells that do not permit long-term culture or completion of the growth cycle. This is reflected in the distinct growth dynamics observed between supporting and non-supporting cell lines, particularly in the differing proportions of bacteria interacting directly with the monolayer versus those remaining unattached in the culture medium (Fig. 2C; Fig. 2D; Supplementary Figure 3). The coexistence of attached and planktonic *T. pallidum* populations may reflect distinct physiological or host-interaction states during *in vitro* growth. The differential persistence of these populations across supportive and non-supportive cell lines suggests that attachment dynamics may influence bacterial survival, replication, or adaptation to the host environment. Future transcriptomic, proteomic, and metabolomic studies will be required to determine whether these populations represent functionally distinct pathogenic states. Consequently, human monolayers that enable long-term maintenance of *T. pallidum* may represent a more appropriate model for studying *T. pallidum*–host interactions than cell lines used only for short-term exposure. In addition, live-cell imaging also demonstrated that *T. pallidum* is capable not only of polar attachment but also of active surface crawling, offering a new perspective on mechanisms underlying tissue invasion and expanding our understanding of *T. pallidum* motility and early host interactions.

Despite our co-culture experiments being limited to four weeks in duration, at the time of writing we have maintained *T. pallidum* in CAL-39 for 7 months and HepG2 for 11 weeks, suggesting that longer-term co-cultivation with human cell lines is feasible and will be important for dissecting additional aspects of *T. pallidum* stability and host-cell adaptation. This shift toward humanized culture systems is further supported by the study from Bosak et al. (22), who demonstrated long-term propagation of eight *T. pallidum* strains using human foreskin fibroblast primary cell lines (HFF1, HFFC, and MoNa), although with slightly reduced replication kinetics relative to rabbit Sf1Ep cells. Because CAL-39 cell line generally supports *T. pallidum* growth in a manner comparable to Sf1Ep cells and outperformed HepG2, we propose it as the more suitable human cell line to replace Sf1Ep cells. Although this study primarily focused on the SS14 strain, reflecting its dominance among contemporary clinical isolates, we also demonstrated that CAL-39 cells support the Nichols strain for several weeks (Supplementary Figure 4). This confirms the utility of the cell line across both major, genetically distinct lineages of *T. pallidum*.

While cell systems cannot fully recapitulate the complexity of infection in the human body, such as tissue architecture, vascularisation, and immune pressure, they offer a reproducible and experimentally accessible platform that increases clinical relevance and enables mechanistic studies of human–pathogen interactions central to syphilis pathogenesis. Compared with more physiologically complex models such as organoids or *ex vivo* explants, which may ultimately provide additional insights but remain technically demanding, the human co-culture systems presented here provide a practical intermediate approach with the added advantage of supporting longer-term culture and potentially capturing previously uncharacterised stages of the bacterial growth cycle. Our discovery will likely enable and guide additional research laboratories that have these and other similar human cell lines already available to adopt the *in vitro* cultivation system for *T. pallidum*. Building on this system, our future work will focus on defining molecular determinants of *T. pallidum* motility, characterising host factors that influence attachment and movement, and dissecting the balance between planktonic and adherent populations.

## Methods

### Human cell lines

Human cell lines to be tested were selected from the Cell Model Passports database (https://cellmodelpassports.sanger.ac.uk/), which contains more than 2000 well-characterised cell lines from diverse tissues. Lines with known tolerance for hypoxia tolerance and slow division rate were prioritised, while the faster-growing HEK-293 cell line was included as a reference (Table 1). Cell lines were also selected to represent anatomically relevant tissue sites associated with *T. pallidum* infection and systemic dissemination.

### Culture conditions and T. pallidum strain

CAL-39, SW954, JEG-3, HepG2, and HEK-293 cells were cultured in filtered, sterile DMEM/F-12 (Gibco 31330-038, ThermoFisher Scientific, UK) supplemented with 10% FBS. JAR cells were cultured in RPMI (Gibco 52400-025, ThermoFisher Scientific, UK) supplemented with 10% FBS and 4.5 mg/mL glucose. Sf1Ep cells were maintained in medium as described by Edmondson et al., 2018 (3). TpCM-2 medium was used without modification for all human cell line co-cultures, and no cell line–specific adaptations were introduced. All cell lines were passaged at confluence (every 3–7 days), and media were changed every 3–5 days depending on cell line and degree of medium acidification. For experimental wells used for *T. pallidum* culture, 50,000 cells were seeded per well in 6-well plates one day prior to inoculation. The *T. pallidum* SS14 strain engineered to express GFP (23) was used for all experiments and was grown *in vitro* as previously described (3) either with standard Sf1Ep cells or with the indicated human cell lines (Table 1). The growth of *T. pallidum* was monitored by dark-field microscopy by assessing both cell counts and motility. *T. pallidum* cell counts were quantified at multiple time points, and mean values with associated variability were plotted over time. Monitoring was performed either at 7-day intervals, as recommended, or daily to generate *T. pallidum* growth curves, with each 24-hour time point analysed from a separate plate that was inoculated simultaneously with an identical inoculum. Growth was assessed in both *T. pallidum* cells attached to the monolayer, following standard culture evaluation procedures (3), and in non-attached, planktonic cells present in the culture medium. For analysis of planktonic cells, 4 mL of culture medium were centrifuged at 10,000 × g for 20 min to pellet *T. pallidum* cells, and the pellet was resuspended in 1 mL of TPCM-2 prior to microscopy. While high-speed centrifugation is likely to disrupt the fragile membrane of *T. pallidum* and reduce cell viability, this was not critical in this context, as the goal was solely to count cells and they were not intended for further sub-passaging.

### Statistical analysis

All statistical analyses and figure generation were performed in R (version 4.5.2). Total *T. pallidum* counts in different cell lines were compared using one-way ANOVA with Tukey’s HSD post hoc test. Temporal *T. pallidum* growth across all cell lines was analysed with two-way ANOVA for the main effects of cell line, day, and their interaction. When significant differences were found, a linear model was used to determine pairwise comparison for each day p-values. *T. pallidum* motility was compared using the Kruskal–Wallis’s test, with Dunn’s post hoc test and Benjamini–Hochberg correction for pairwise comparisons. The relationship between mammalian cell division time and *T. pallidum* yield (log_10_ fold change) was assessed by linear regression. *T. pallidum* in planktonic state in the media was compared between supporting and non-supporting cell lines using Wilcoxon rank-sum tests performed separately for each day.

Each measurement was performed using three biological replicates. Although this limited statistical power for some comparisons, the number of replicates was dictated by experimental complexity and the need to complete treponemal counting and motility assessments within a narrow time window. Increasing the number of replicates could have been detrimental to data reliability, as treponemes do not maintain viability indefinitely after harvest.

### Live cell imaging

For live-cell imaging, a Leica THUNDER Imager (Leica Microsystems, Milton Keynes, UK) was used, selected for its computational background removal and low phototoxicity. To maintain appropriate culture conditions during imaging, a cage incubator, an on-stage gas-tight chamber, and gas and temperature controllers (Okolab, Rovereto, Italy) were employed. These were connected to a premixed gas cylinder (1.5% O2, 5% CO2, balance N2; BOC, Cambridge, UK) to provide a microaerophilic atmosphere suitable for *T. pallidum* growth and to maintain the temperature at 34 °C. *T. pallidum* was co-cultured with different cell lines in microscopy dishes compatible with the gas-tight chamber (Nunc™ Glass Bottom Dishes, cat. no. 150680, Thermo Fisher Scientific, UK). A 63× oil-immersion objective was used to monitor *T. pallidum* motility and interactions. Motility phenotypes were assessed qualitatively by visual inspection of live-cell imaging data; no formal tracking-based quantification of speed, directionality, or persistence was performed.

### Scanning and transmission electron microscopy (SEM and TEM)

*T. pallidum* was cultured under standard conditions for 7 days in the presence of monolayers of various mammalian cells grown on sterile circular coverslips (12 mm; Paul Marienfeld GmbH & Co. KG, Lauda-Königshofen, Germany). Coverslips were placed one per well in 24-well plates and used for SEM or TEM sample preparation. Immediately after removal of the plates from the incubator, cells were inactivated and fixed by exchanging the culture medium for 2% glutaraldehyde and 4% paraformaldehyde (PFA) in 0.1M sodium cacodylate (pH 7.42) for 2 hours (all steps at room temperature). They were then rinsed in cacodylate buffer (3 x 10 minutes), post-fixed in cacodylate buffered 1% osmium tetroxide for a further 2 hours, rinsed in buffer again and mordanted in either 1% buffered tannic acid (TEM) or 1% thiocarbohydrazide (SEM) for 1 hour. Then, for TEM, cells were washed in sodium sulphate for 10 minutes, rinsed 6 x 2 minutes in distilled water, dehydrated in an ethanol series, stained *en bloc* in 1% uranyl acetate at the 20% stage, embedded using an Epoxy resin embedding kit (Epoxy embedding medium 45345, Sigma-Aldrich – Merck, Gilingham, UK), and cured in an oven at 60°C for 48 hours. Areas containing cells were cut from the well and ultrathin-sectioned at 60nm on a Leica UC6 ultramicrotome. Sections were collected onto EM grids, contrasted again with uranyl acetate and lead citrate and analysed on a 120kV Tecnai Spirit Biotwin TEM with a Tietz F4.16 CCD. For SEM, samples were further impregnated with osmium tetroxide for 1 hour, rinsed in buffer for 3 x 10 minutes and distilled water for 6 x 2 minutes, dehydrated in an ethanol series, the coverslips removed and critical point dried in a Leica EM CPD300, mounted onto SEM stubs with conducting paste, sputter-coated with 5nm of gold in a Leica ACE600 sputter-coater and analysed on a Hitachi SU8030 SEM.

## Figures

**Video file link (real speed *T. pallidum* interaction with the host cell)**

https://drive.google.com/file/d/14sPVNNl79a9wy-LJo6AXBiy3GagYyBuB/view?usp=drive_link

## Acknowledgments

This work was supported by the Biotechnology and Biological Sciences Research Council (BBSRC) (grant number BB/010589), the Bill & Melinda Gates Foundation (INV-051483), and the Wellcome funding (grant 220540/Z/20/A). We also thank Dr. Matthew Garnett (Wellcome Sanger Institute, Wellcome Genome Campus, Hinxton, UK) for providing access to cell lines in the Cell Model Passports dataset.

## Supplementary material

**Supplementary Figure 1.**
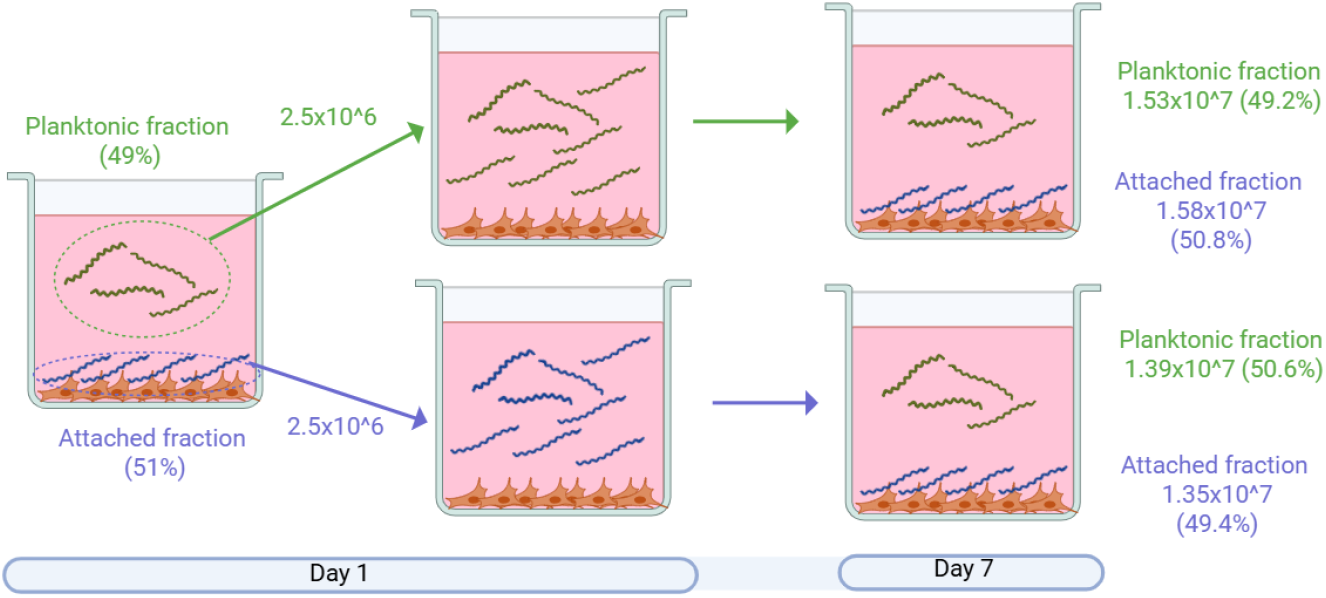
Schematic representation of re-seeding experiments demonstrating that *T. pallidum* cells recovered exclusively from culture supernatant can reattach to fresh host cell monolayers and complete a full replication cycle. Equal numbers of *T. pallidum* (2.5 × 10^6^), either collected from the planktonic fraction (culture medium only) or from the attached fraction (host cell–associated bacteria released by trypsinization), were inoculated into fresh media and monolayers. After 7 days, both conditions yielded comparable total bacterial counts and similar proportions of planktonic and host cell–associated fractions. These results demonstrate that planktonic *T. pallidum* remain competent for reattachment and replication. Values represent mean counts and percentages from three independent biological replicates.

**Supplementary Figure 2.**
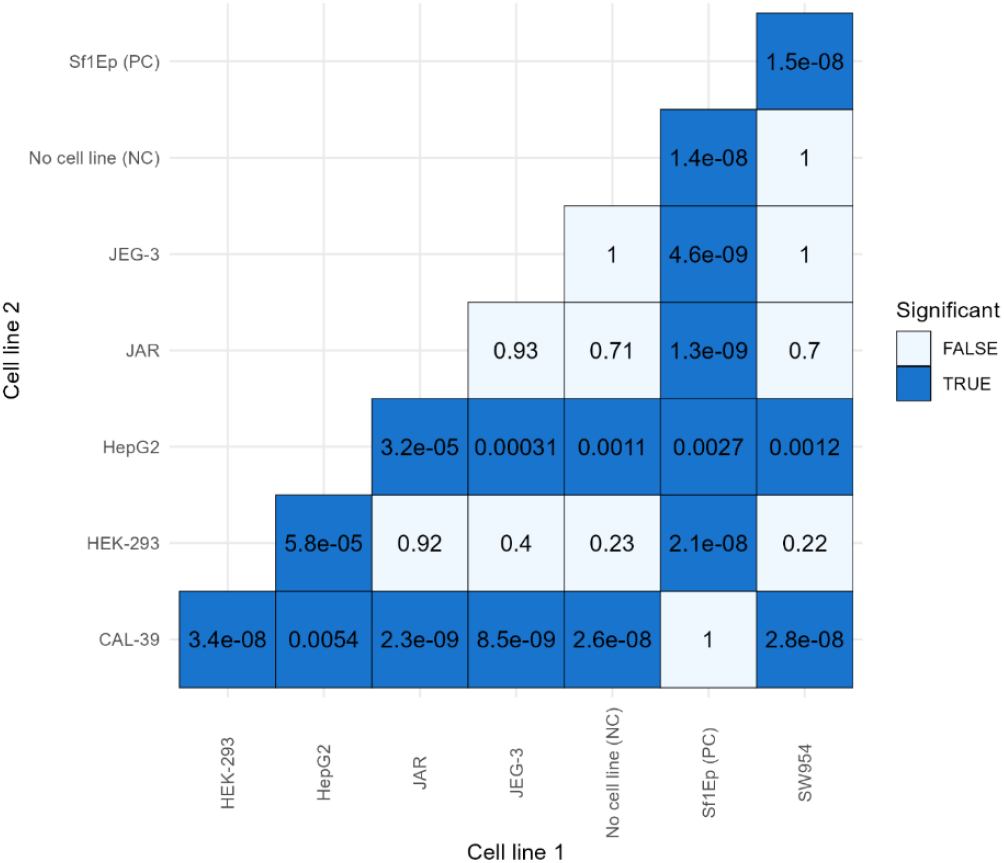
Triangular heatmap of pairwise TukeyHSD post hoc comparisons of *T. pallidum* counts between cell lines. Each cell line shows the adjusted p-value for a given pairwise comparison; blue tiles indicate statistically significant differences (p < 0.05).

**Supplementary Figure 3.**
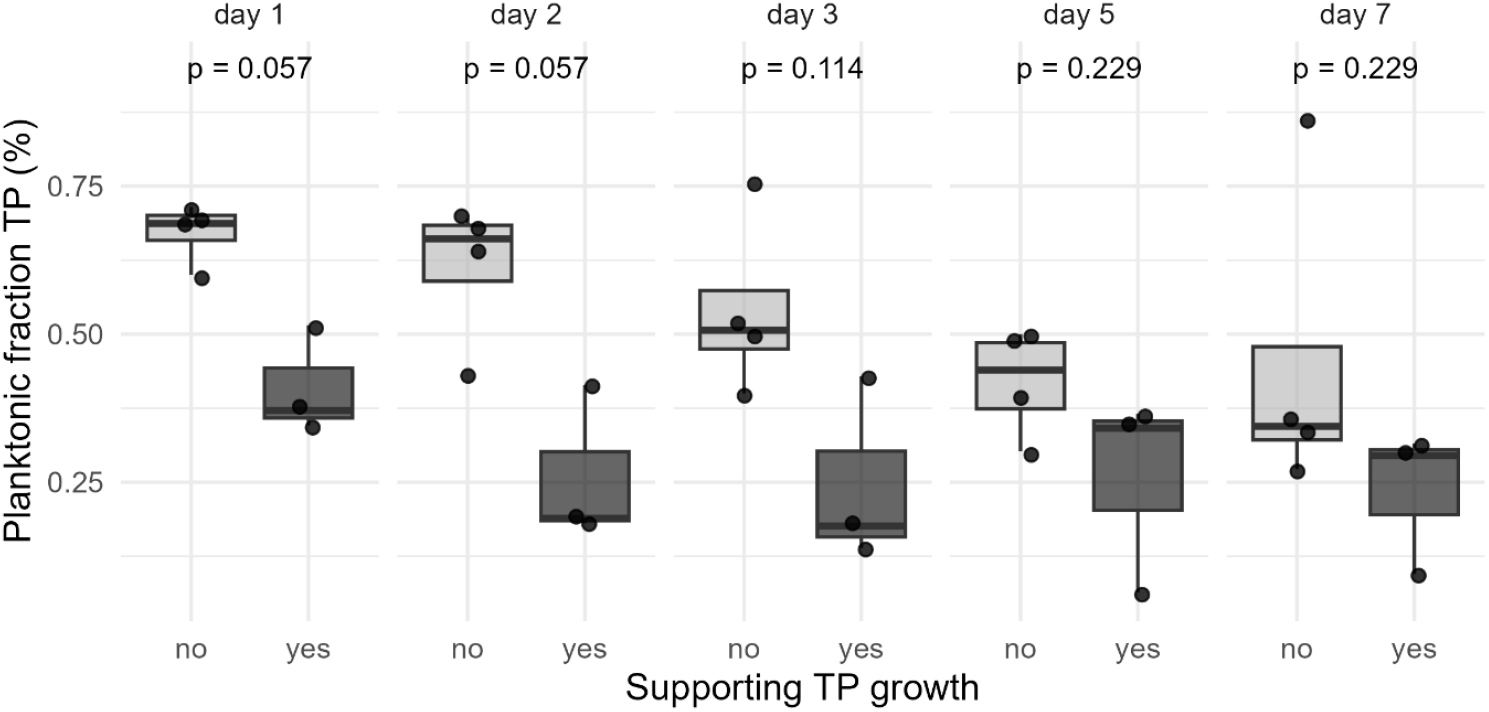
Planktonic fraction of *T. pallidum* in supporting (Sf1Ep, CAL-39, HEPG2) and non-supporting (HEK-293, SW954, JAR, JEG-3) cell lines across days 1, 2, 3, 5, and 7. Boxplots show the distribution of planktonic *T. pallidum* fractions for each group, with individual data points overlaid. Wilcoxon rank-sum tests were performed separately for each day (day 1, p = 0.057; day 2, p = 0.057; day 3, p = 0.114; day 5, p = 0.229; day 7, p = 0.229).

**Supplementary Figure 4.**
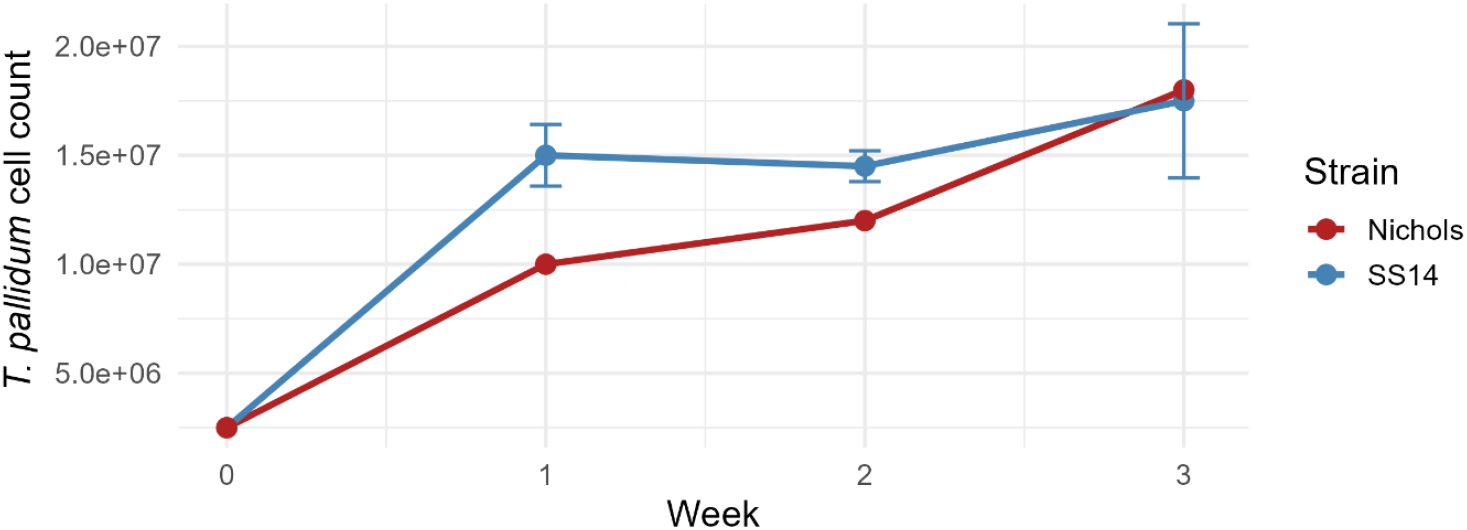
Growth of *T. pallidum* SS14 and Nichols in the presence of CAL-39. A linear mixed-effects model showed no significant strain × week interaction (F = 0.0076), indicating no detectable differences in growth trends between SS14 and Nichols.

**Supplementary table 1.** Pairwise contrasts from a linear model comparing CAL-39, HepG2, and Sf1Ep across 10 days. Reported are estimated differences, standard errors, degrees of freedom, t-ratios, adjusted p-values. Significant differences after adjustment were observed primarily at later time points, including CAL-39 vs HepG2 on days 6 and 8, and contrasts involving Sf1EP on days 9 and 10, indicating time-dependent divergence between cell lines.

| Contrast | Day | Difference | Se | Df | T_Ratio | P_Adjusted |
| --- | --- | --- | --- | --- | --- | --- |
| CAL-39 - HEPG2 | day_1 | 3746000 | 5254010 | 58 | 0.712979 | 0.756833 |
| CAL-39 - SF1EP | day_1 | 2313333 | 5254010 | 58 | 0.440299 | 0.898848 |
| HEPG2 - SF1EP | day_1 | -1432667 | 5254010 | 58 | -0.27268 | 0.959869 |
| CAL-39 - HEPG2 | day_2 | 9020667 | 5254010 | 58 | 1.716911 | 0.207578 |
| CAL-39 - SF1EP | day_2 | 6734000 | 5254010 | 58 | 1.281688 | 0.411135 |
| HEPG2 - SF1EP | day_2 | -2286667 | 5254010 | 58 | -0.43522 | 0.901043 |
| CAL-39 - HEPG2 | day_3 | 10901333 | 5254010 | 58 | 2.07486 | 0.104008 |
| CAL-39 - SF1EP | day_3 | 7410000 | 5254010 | 58 | 1.410351 | 0.342288 |
| HEPG2 - SF1EP | day_3 | -3491333 | 5254010 | 58 | -0.66451 | 0.784895 |
| CAL-39 - HEPG2 | day_4 | 8213333 | 5254010 | 58 | 1.563251 | 0.269663 |
| CAL-39 - SF1EP | day_4 | 5490000 | 5254010 | 58 | 1.044916 | 0.551868 |
| HEPG2 - SF1EP | day_4 | -2723333 | 5254010 | 58 | -0.51833 | 0.862722 |
| CAL-39 - HEPG2 | day_5 | 6898667 | 5254010 | 58 | 1.313029 | 0.393771 |
| CAL-39 - SF1EP | day_5 | 3283333 | 5254010 | 58 | 0.62492 | 0.80708 |
| HEPG2 - SF1EP | day_5 | -3615333 | 5254010 | 58 | -0.68811 | 0.771345 |
| CAL-39 - HEPG2 | day_6 | 13632667 | 5254010 | 58 | 2.594717 | 0.031641 |
| CAL-39 - SF1EP | day_6 | 2283333 | 5254010 | 58 | 0.434589 | 0.901316 |
| HEPG2 - SF1EP | day_6 | -1.1E+07 | 5254010 | 58 | -2.16013 | 0.086819 |
| CAL-39 - HEPG2 | day_7 | 11850000 | 5254010 | 58 | 2.25542 | 0.070455 |
| CAL-39 - SF1EP | day_7 | 2673333 | 5254010 | 58 | 0.508818 | 0.867355 |
| HEPG2 - SF1EP | day_7 | -9176667 | 5254010 | 58 | -1.7466 | 0.196854 |
| CAL-39 - HEPG2 | day_8 | 16310000 | 5254010 | 58 | 3.104296 | 0.00817 |
| CAL-39 - SF1EP | day_8 | 4830000 | 5254010 | 58 | 0.919298 | 0.630397 |
| HEPG2 - SF1EP | day_8 | -1.1E+07 | 5254010 | 58 | -2.185 | 0.082271 |
| CAL-39 - HEPG2 | day_9 | 5723333 | 5254010 | 58 | 1.089327 | 0.524476 |
| CAL-39 - SF1EP | day_9 | -1.7E+07 | 5254010 | 58 | -3.32698 | 0.004297 |
| HEPG2 - SF1EP | day_9 | -2.3E+07 | 5254010 | 58 | -4.41631 | 0.00013 |
| CAL-39 - HEPG2 | day_10 | 2703333 | 5254010 | 58 | 0.514528 | 0.864582 |
| CAL-39 - SF1EP | day_10 | -2.8E+07 | 5254010 | 58 | -5.41935 | 3.58E-06 |
| HEPG2 - SF1EP | day_10 | -3.1E+07 | 5254010 | 58 | -5.93388 | 5.22E-07 |

